# An iron-sulfur cluster as a new metal centre in a flavodiiron protein

**DOI:** 10.1101/2024.04.03.587993

**Authors:** Maria C. Martins, Célia M. Silveira, Leonor Morgado, Miguel Teixeira, Filipe Folgosa

**Affiliations:** Instituto de Tecnologia Química e Biológica António Xavier, Universidade Nova de Lisboa, Av. da República, 2780-157 Oeiras, Portugal; Associate Laboratory i4HB–Institute for Health and Bioeconomy, and UCIBIO Applied Molecular Biosciences Unit, Chemistry Department, NOVA School of Science and Technology, Universidade NOVA de Lisboa, Largo da Torre, 2829-516 Caparica, Portugal

**Keywords:** FeS cluster, ferredoxin, flavodiiron protein, *Syntrophomonas*, flavin, diiron

## Abstract

*Syntrophomonas wolfei* contains two distinct multiple domain flavodiiron proteins (FDPs), of Classes H and E, presumably acting as oxygen reductases to protect this anaerobic bacterium from oxidative stress due to exposure to environments containing, even if only transiently, oxygen. The Class E FDP was predicted to have, besides the two core domains characteristic of this type of enzymes, an extra C- terminal domain putatively harbouring an iron-sulfur centre. Bioinformatic analyses showed that, thus far, Class E FDPs are only present in three other bacteria of the *Syntrophomonas* genus: *Syntrophomonas palmitatica*, *Syntrophomonas zenhnderi* and *Thermosyntropha lipolytica.* In this work, we extensively characterized the enzyme from *Syntrophomonas wolfei* (wild type, site directed mutants and truncated domains) and showed unequivocally, using EPR and Resonance Raman spectroscopies, that indeed it contains a [3Fe- 4S]^1+/0^ centre, a novelty in the field of FDPs. Structure prediction using Alphafold indicated some similarities to [3Fe4S]^1+/0^ containing ferredoxins. The reduction potentials of each cofactor were determined: +70 mV, -5/-70 mV and -90 mV for the FeS, diiron centre and flavin, respectively.

## Introduction

Flavodiiron proteins (FDPs) constitute an extensive family of metalloenzymes present in all life domains with crucial roles in oxygen and Reactive Oxygen Species (ROS, namely H_2_O_2_) detoxification and/or NO detoxification, through the reduction of these species either to H_2_O or N_2_O, respectively. [1,2]

All FDPs are composed by a minimal catalytic unit: a dimer (mostly a homodimer), in which each monomer has two domains: a metallo-β- lactamase-like domain, harbouring the catalytic diiron site and a flavodoxin-like domain with an FMN moiety. The majority of the flavodiiron proteins already characterized have only this core, but many FDPs were found by bioinformatics and structure prediction studies to have additional structural domains after the flavodoxin-like domain, namely those from Clostridia and protozoa such as *Trichomonas vaginalis.* FDPs were classified in nine classes on the basis of their domain’s architecture, containing additional cofactors: for example, rubredoxins and NADH:rubredoxin oxidoreductases-like domains [1,3]. For the first time, in the present work, we describe an FeS cluster domain fused with the catalytic core of an FDP: a representative from class E FDPs (FDP_E). This class contains a C-terminal domain identified as the protein domain family “Fer4_19” (Pfam06902 protein family), that may harbour either a [3Fe-4S]^1+/0^ or a [4Fe-4S]^2+/1+^ centre.

FeS clusters are the most ubiquitous metalloproteins, present in the three life kingdoms and are involved in several key biological processes like photosynthesis, respiration and gene regulation. These proteins possess a wide range of reduction potentials and diverse structural motifs which allow them to interact with several redox partners and consequently work as electron carriers in a multiplicity of biological processes. Besides its most well characterized function, as electron carriers, FeS clusters constitute part of the active sites of several enzymes and are involved in functions such as initiation/stabilization of radical chain reactions and reduction of disulfide bonds. Moreover, FeS clusters are key players in cellular response mechanisms against oxidative and metal stresses, due to the frequent sensitivity of these centres to an oxidative environment and their wide range of redox states allows them to act as cell sensors [4–7].

FeS centres can be classified in different groups on the basis of the number of iron and sulfur atoms, structural motifs and spectroscopic and electrochemical properties. The most common FeS centres are organized in four main groups: rubredoxins ([1Fe−4S]), ferredoxins with [2Fe−2S]^2+/1+^, [3Fe−4S]^1+/0^ or [4Fe−4S]^2+/1+^ centres, Rieske proteins, and high-potential iron−sulfur proteins (HiPIPs, 4Fe-4S]^3+/2+^) [5,8].

Here, we present a detailed description of the biochemical and spectroscopic features of the three domains of the class E FDP from *Syntrophomonas wolfei*, with a special focus on the characterization of the iron-sulfur centre present in the C-terminal domain, a novelty in the field of FDPs. For this purpose, we analysed the FDP_E holo-enzyme, a truncated form containing only the C-terminal domain (FDP_E_Cter) and two FDP_E site-directed mutants. In these mutants, a histidine residue present in the conserved motif characteristic of the “Fer4_19” family and eventually involved in iron coordination of the FeS cluster, was replaced by an alanine (E_H441A) and by a cysteine (E_H441C) in order to assess the putative role of this conserved histidine in the iron coordination of the FeS cluster.

## Results and Discussion

### Amino acid sequence analysis and structural predictions

The Swol_2363 gene, from *Syntrophomonas wolfei subsp*. *Wolfei* str. Goettingen G311, encodes for a class E FDP, with 508 amino acids per monomer. The first 400 residues comprise the flavodiiron core, divided in the metallo-β-lactamase domain, in which the ligands for the diiron site are fully conserved (H81, E83, D85, H86, H149, D168, and H229, FDP_E numbering used in this article). Amino acid sequence alignment of FDP_E flavodiiron core with the core domains from previously characterized FDPs, showed an identity of circa 30%. When compared with the core domain of the multidomain FDP_H [9], also from *S. wolfei*, the identity increases to 63%.

Besides the flavodiiron core, FDP_E has an extra C-terminal domain characteristic and exclusive of this class of FDPs, containing a putative FeS cluster. This FeS domain has homology with a protein domain family “Fer4_19” (Pfam06902) which is characterized by the presence of a conserved motif (CXHX_3_CX_32_CP). The presence of three strictly conserved cysteines indicate that this domain may harbour a [3Fe- 4S]^1+/0^ or even a [4Fe-4S]^2+/1+^ cluster considering that FeS centres can possess non-conventional ligands, such as histidines, that may act as the fourth ligand [10]. Interestingly, one of the residues of this conserved motif is a histidine. The “Fer4_19” is part of the “Fer4_9” superfamily (Interpro IPR017896) which is described as a [4Fe-4S]^2+/1+^ ferredoxin-type, iron-sulfur binding domain containing family of proteins. The pattern of cysteine residues in the iron-sulfur region is sufficient to detect this class of [4Fe-4S]^2+/1+^ binding proteins (https://www.ebi.ac.uk/interpro/entry/InterPro/IPR017896/).However, none of these proteins has yet been characterized.

AlphaFold [11,12] structure prediction was used to build the model of the full-length protein (Figure 1, panel A), highlighting the presence of the three predicted domains with an extensive loop (20 amino acids) connecting the FDP-core to the FeS cluster domain. Additionally, we generated the AlphaFold model for the metallo-β-lactamase-like (residues 1 to 243) and flavodoxin-like (residues 251 to 398) domains and the flavodiiron core of FDP_E (residues 1 to 398). The core model was used to perform a structural alignment with the crystallographic structure of *E. coli* FDP (PDB code: 4D02), obtaining a poor alignment with a considerably high rmsd value of 6.38Å (256 atoms) [13]. By analysing the structural superposition, the high rmsd value between the two structures was caused by a different relative orientation between the two core domains (as observed also for Class F from *Thricomonas vaginalis* [14] and predicted for the class H from *Syntrophomonas wolfei* [9]) (Figure 1, panel B). This feature may be related with two different factors. The first and simpler, the prediction of a longer linker between the metallo-β-lactamase and the flavodoxin domains, which is present in FDP_E and absent in the class A and B FDPs, such as the *E. coli* FDP structure (5 amino acids in *E. coli* FDP versus 9 amino acids in FDP_E), may allow some degree of flexibility and lead to the different relative orientation. The second hypothesis may be related with the charge surface in-between the two core domains. By analysing the crystallographic structures available for the different FDPs, most have either opposite or close to neutral charges in-between the surfaces of the two core-domains structure, contrary to what was predicted for the class E FDP model. In agreement with this observation, lower rmsd values were obtained, 1.23 and 2.82 Å for the metallo-β-lactamase and flavodoxin domains, respectively, when the models of the two core domains were superimposed separately (Figure 1, panel C).

**Figure 1.**
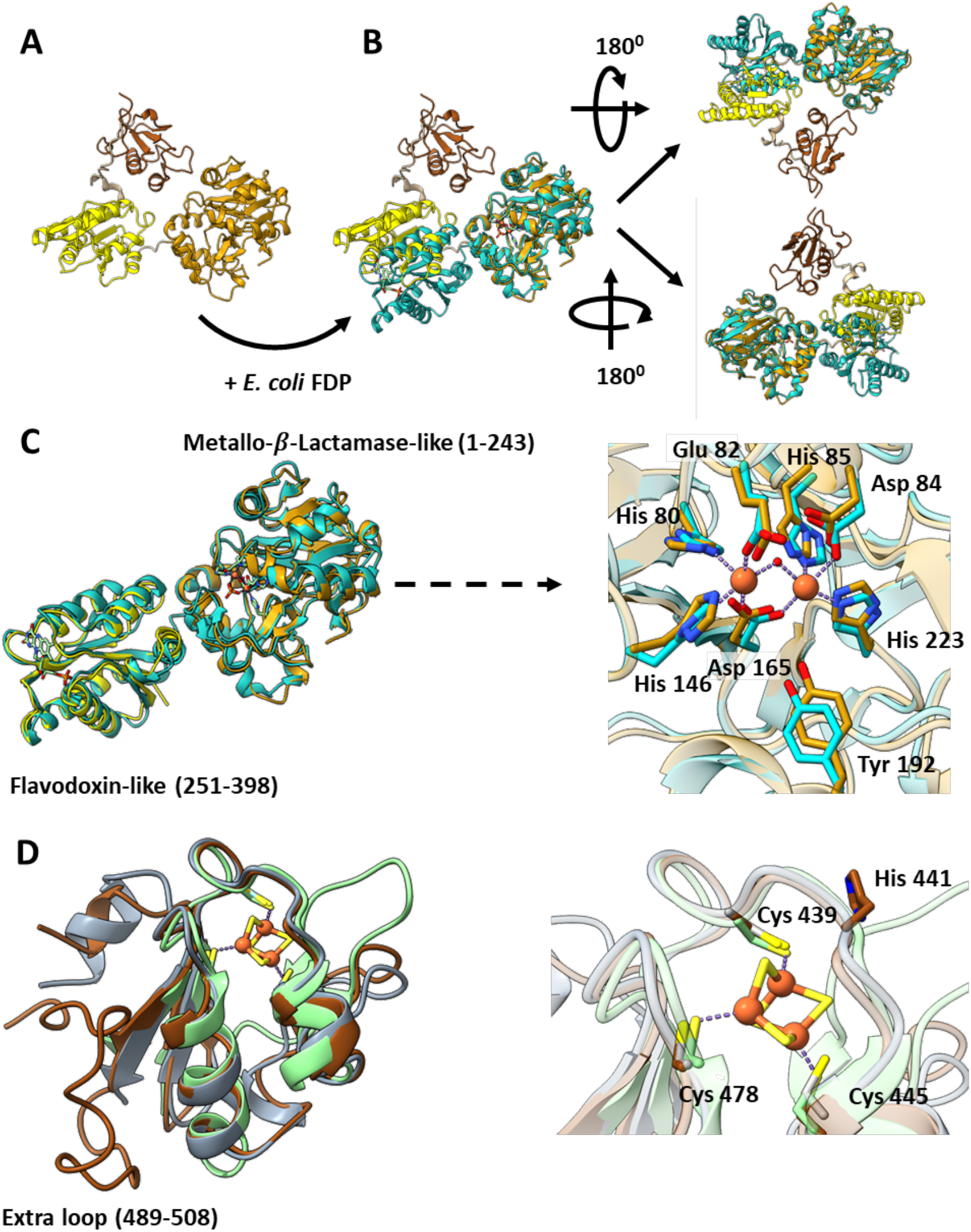
Structure prediction using AlphaFold. **(A).** FDP_E, highlighting the three domains in distinct colors: orange - metallo-β-lactamase domain; yellow – flavodoxin domain; red – FeS domain **(B).** Structure superposition of the two predicted core domains of *S. wolfei* FDP_E with the crystallographic structure of *E. coli* FDP core (PDB 4D02, blue) in two different spatial orientations. **(C).** Structure superposition of the two predicted core domains (superimposed separately) of *S. wolfei* FDP_E with the corresponding crystallographic structure of *E. coli* FDP domains (PDB 4D02, blue). **(D).** Structure superimposition of the FDP_E_Cter (residues 417-508 of FDP_E, brown colour) with the crystallographic structure of *Mycobacterium tuberculosis* (PDB code: 8AMP, green) and the AlphaFold model of YjdI from *E. coli* (Uniprot P0AF59, grey).

As this is the first time such type of C-terminal FeS cluster domain is characterized in FDPs and there are no structural studies of other proteins belonging to the “Fer4_19” family, we searched the literature for FeS proteins with similar structure arrangements and a similar number of residues (c.a. 100 residues), namely recurring to the Protein Fold Recognition Server tool [15]. Using this method, we retrieved a number of different proteins, but the top hits found corresponded to small ferredoxins.[5,16] By performing structural alignments between the AlphaFold model of FDP_E_Cter (residues 417-508 of FDP_E) with different classes of ferredoxins with determined crystallographic structures, we concluded that this domain shows a higher structural homology with [3Fe-4S]^1+/0^ ferredoxins, namely with the one from *Mycobacterium tuberculosis* (PDB code: 8AMP) with a rmsd value of 4.6 Å (60 matching amino acids) (Figure 1 panel D) [17]. The AlphaFold model of the FDP_E_Cter was also compared with the Alphafold model of YjdI from *E. coli* (Uniprot P0AF59), which is part of the “fer4_19” family and was already spectroscopically characterized [18]. The structural alignment between these two models retrieved a rmsd value of 1.5 Å (72 matching amino acids), showing a large similarity between these two models and, eventually proteins. Interestingly, both these models present a section, with c.a. 12 amino acids, in the N-terminal side that is not present in the *Mycobacterium tuberculosis* ferredoxin. Amino acid sequence alignments of FDP_E_Cter with the sequences of YjdI from *E. coli, Mycobacterium tuberculosis* ferredoxin and other [3Fe- 4S]^1+/0^ ferredoxins (Figure S1) showed pairwise identities ranging between 15 and 30% (for *Rhodopseudomonas palustris* HaA2 ferredoxin and YjdI from *E. coli*, respectively). It also showed that the FeS cluster coordinating cysteines are conserved among them. The exception, as was already observed in the structural superposition, was the presence of a 20 amino acids extension in the C-terminal side, that is putatively responsible for the presence of an extra loop in the structure.

The class E FDP is so far phylogenetically restricted, since it was only found in genomes of organisms from the Syntrophomonadaceae family, specifically in *Syntrophomonas wolfei*, *Syntrophomonas palmitatica*, *Syntrophomonas zehenderi* and *Thermosyntropha lipolytica* (as of March 2024). The FDP E sequences of these organisms present an identity of 55%.

### General biochemical properties of FDP_E

FDP_E (full-length protein) and the truncated construct FDP_E_Cter, corresponding to the residues 401 to 508, were successfully produced in *E. coli* and purified to homogeneity, under anaerobic conditions. The flavin content of FDP_E was evaluated by UV-Visible spectroscopy and determined as 27% of flavin/per protein monomer. The iron incorporation was determined by ICP for FDP_E and FDP_E_Cter, obtaining 26% and 37% iron/protein, respectively.

The quaternary structure of the two constructs in solution were determined by size exclusion chromatography. Both proteins were isolated as dimers, with molecular masses of 103.8 kDa and 18.5 kDa for the full length and C-term, respectively, consistent with the calculated mass for each monomer based on the amino acid sequences, 57.5 kDa for FDP_E and 12 kDa for FDP_E_Cter. Until now, all the FDPs already characterized exist in solution as dimers or tetramers (dimer of dimers), a configuration that is crucial for an efficient intramolecular electron transfer between the FMN from one monomer (the primary electron acceptor of the core enzyme) and the diiron catalytic site of the other monomer [19].

The UV-Visible spectrum of as purified FDP_E (Figure 2), is dominated by the characteristic flavin features of the flavodoxin domain, with maxima at 360 and 450 nm, and as expected, the diiron centre, due to its low absorptivity, is not detected. Moreover, the contribution of the FeS cluster is indistinguishable from the flavin contribution since [3Fe- 4S]^1+/0^ or [4Fe-4S]^2+/1+^ centres are characterized by a broad absorbance band with a maximum around 400 nm[5]. In Figure 2, the spectrum of FDP_E_Cter, due to the absence of FMN in this construct, reveals the presence of the iron-sulfur cluster, which exhibits a broad band with a maximum at 410 nm. The spectrum is quite similar to the one of YjdI from *E. coli* [18].

**Figure 2.**
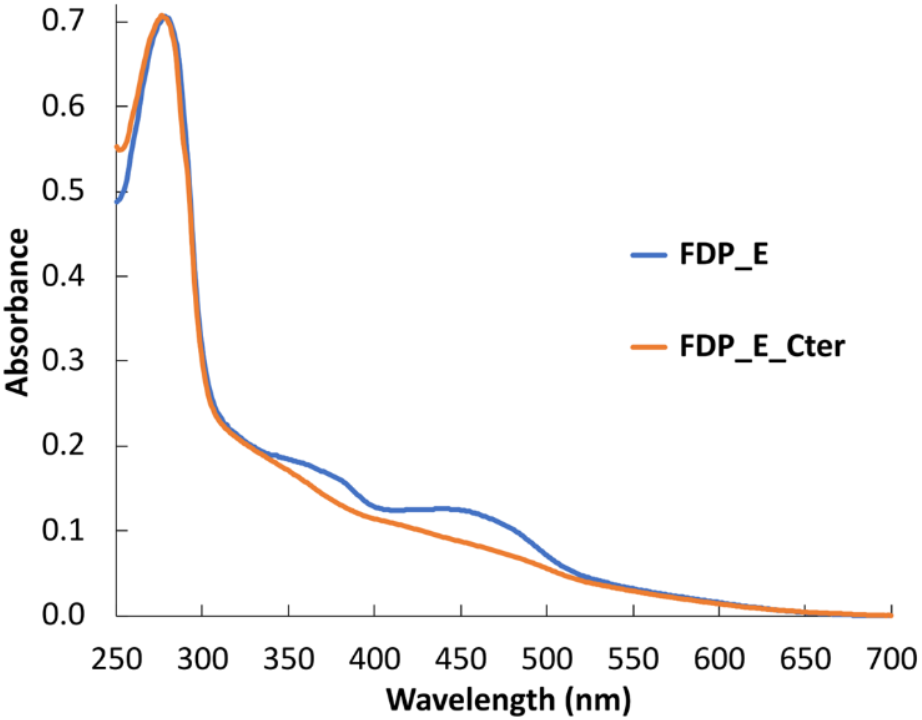
**UV-Visible spectra of 18 μM FDP_E** (blue) **and 30 μM FDP_E_Cter** (orange) in 50 mM Tris-HCl pH 7.5 containing 18 % glycerol.

In Figure S2, are also depicted the UV-Visible spectra of the E_H441A and E_H441C variants of FDP_E, becoming clear that the point mutations of the histidine, localized in the conserved motif characteristic of this FeS domain, did not originate any changes in the electronic spectra. When FDP_E, E_H441A and E_H441C spectra were compared slight differences were observed, which is most likely related with the different flavin content amongst the samples (approximately 6% less flavin incorporation when compared with the wild type protein).

### Metallo-β-lactamase-like domain spectroscopic characterization

Anaerobic redox titration of FDP_E, followed by EPR, was used to determine the parameters of the catalytic diiron centre in its mixed-valence state as well as its reduction potentials. By spectral simulation of the FDP_E sample prepared at -71 mV (Figure 3, panel A), we obtained two g values for the rhombic signal, 1.85 and 1.77. The g_max_ was not observed because of the high contribution of the radical species at g = 2, that distorted the mixed-valence diiron signal. Therefore, for simulation purposes we assumed g = 1.96 for the g_max_ (a typical value of this component in the other diiron centres of FDPs already characterized). The g-values obtained for the diiron centre of FDP_E are slightly higher when compared with the diiron centres of other FDPs, namely *E. coli*, *E. histolytica*, *G. intestinalis* and *T. vaginalis* FDPs [14,20–22].

**Figure 3.**
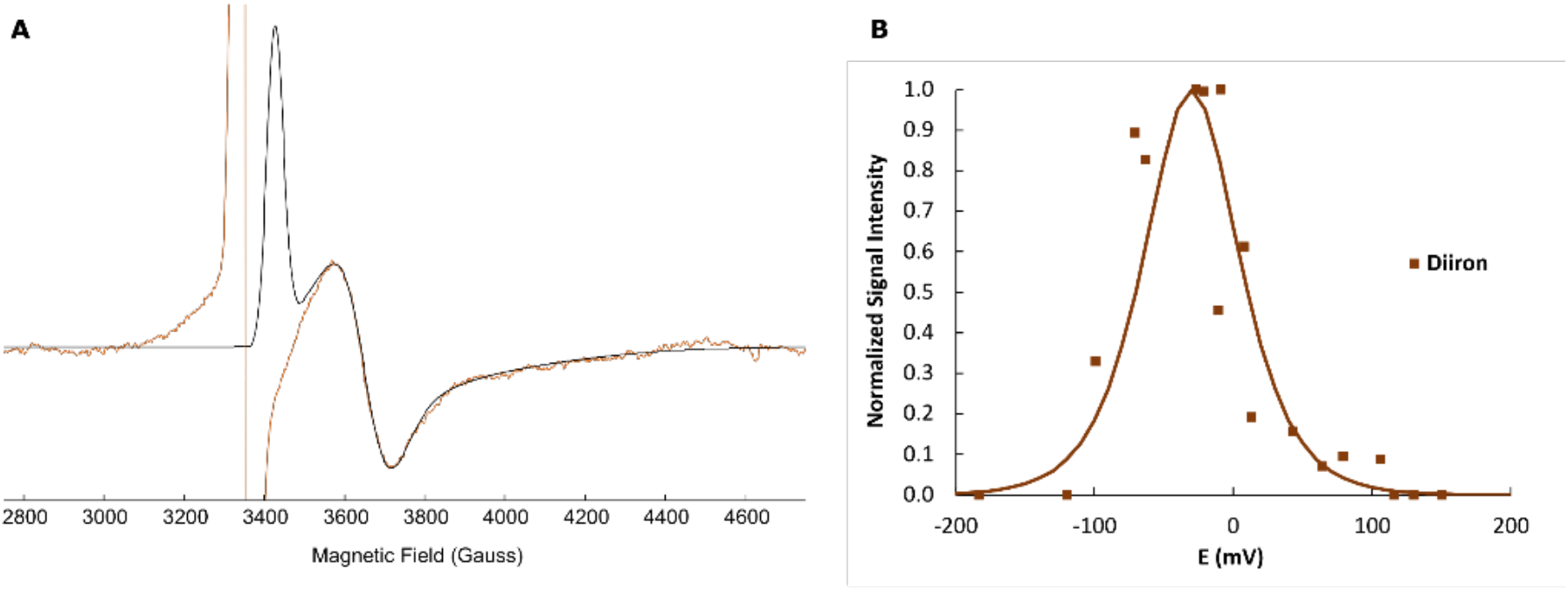
Spectroscopic characterization of the diiron centre of FDP_E. **(A)** EPR spectrum of 150 μM FDP_E at -71 mV (orange). The black line corresponds to the simulated spectrum obtained for the diiron centre using g values of 1.96, 1.85 and 1.77. **(B).** Redox titration monitored by EPR redox following the intensity of the signal at g=1.85. Temperature, 7 K; Microwave frequency, 9.39 GHz; Modulation amplitude, 1.0 mT; Microwave power, 2 mW. Spectral simulations and image preparation were performed using SpinCount [23].

To determine the reduction potentials of the diiron centre we measured the intensity of the resonance at g=1.85 as a function of the different redox potentials. To these experimental points, a Nernst equation for two consecutive monoelectronic transitions was manually adjusted, and reduction potentials of –5 mV and -60 mV (Figure 3, panel B) were determined for each transition. These values are in the same range of those previously determined for the diiron centre of *E. coli* FDP ( -20 and -90 mV) [20] but significantly lower when compared with the diiron centres of *E. histolytica* (+170 and +132 mV)[21], *G. intestinalis* (+163 and +2 mV) [22]and *T. vaginalis* (+190 and +50 mV)[14] FDPs.

### Flavodoxin-like domain spectroscopic characterization

The flavodoxin-like domain was characterized by UV-Visible and fluorescence spectroscopies and reverse phase HPLC. The flavin component of FDP_E dominates the spectrum of the as purified (oxidized) sample (Figure 2) as mentioned above. Upon reduction of FDP_E with an excess of dithionite (even upon overnight incubation) it was not possible to obtain a complete reduction of the flavin, contrary to other FDPs [9,24,25]. The resulting spectrum is very similar to a red (anionic) semiquinone (grey spectrum in Figure 4 panel A), with a drop in the absorbance at 455 nm accompanied by an increase at ≈ 490 and ≈ 385 nm, as was previously obtained in the partial reduction of other FDPs. [25] This effect was also observed during the redox titration where the final spectrum obtained pointed to the semiquinone as the final species, impairing the determination of the second reduction potential (from the semiquinone to the fully reduced species). Therefore, the only reduction potential determined, -90 mV (monitored at 475 nm), refers to the reduction of the oxidized flavin to the semiquinone form (Figure 4 panel B).

**Figure 4.**
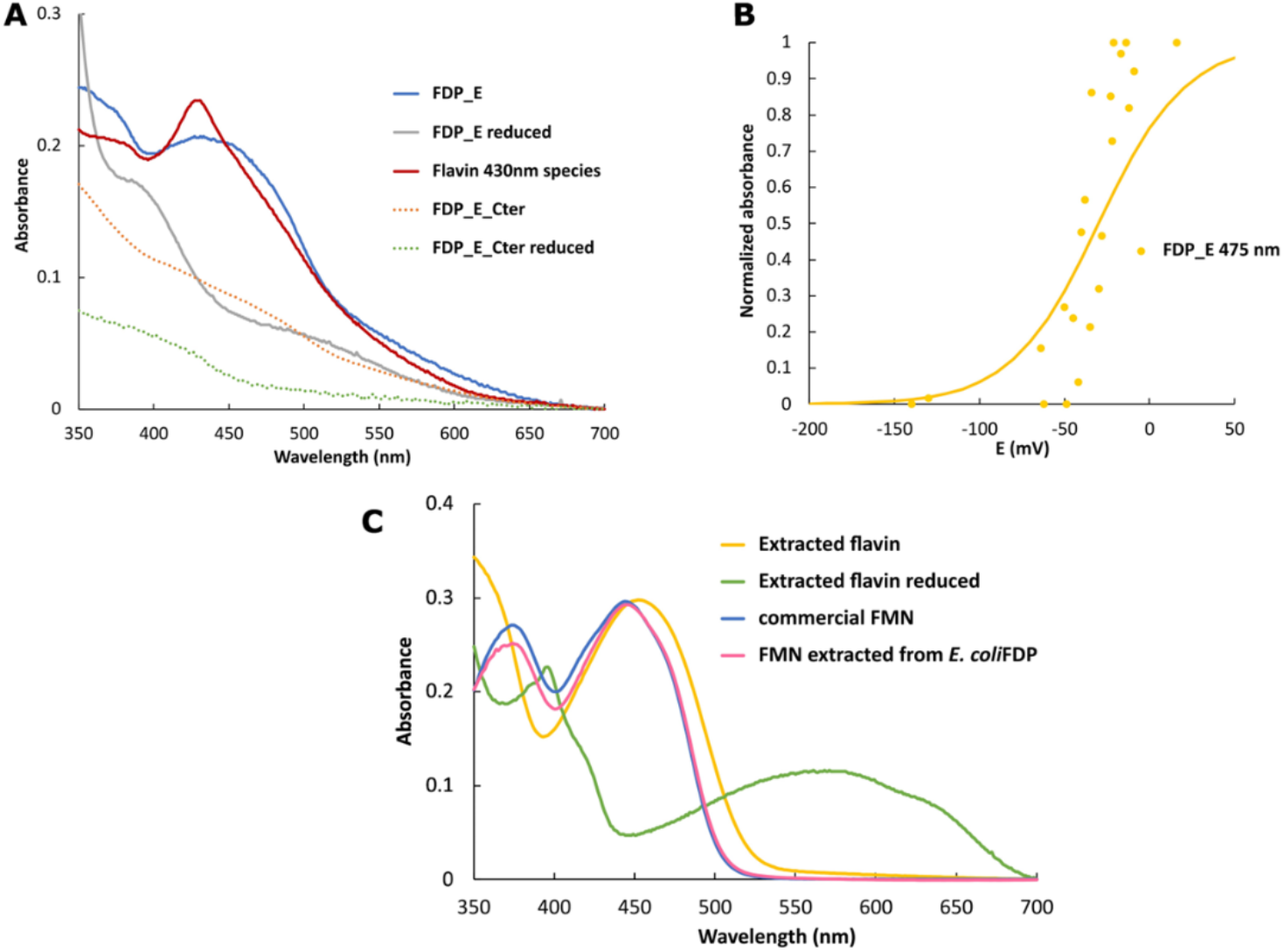
Spectroscopic characterization of the FDP_E flavin. **(A)** UV-Visible spectra of *S. wolfei* FDP_E and FDP_E_Cter, between 350 and 700 nm, in the oxidized form (blue and orange-dotted lines, respectively) and after reduction with sodium dithionite (grey line for FDP_E reduced and green-dotted line for FDP_E_Cter reduced). The red spectrum represents the flavin 430 nm species which arises from the partial reduction of FDP_E. **(B)** Redox titration followed by UV-Visible spectroscopy of FDP_E. In the figure are represented the normalized intensities measured at 475 nm. The titration was performed with 15 µM of protein in 50mM Tris-HCl pH 7.5 containing 18% glycerol. The solid lines correspond to theoretical data using the appropriate Nernst equations as explained in the text. **(C)** UV-Visible spectra of the cofactor isolated from FDP_E as isolated (yellow) and in the reduced form (green spectrum) in comparison with the spectrum of commercial FMN (blue) and the FMN extracted from *E. coli* FDP.

Moreover, in both these experiments, the formation of an unidentified intermediary species was observed, with a maximum at 430 nm that appears upon the addition of substoichiometric amounts of dithionite and at around –12 mV during the redox titration (red spectrum in Figure 4 panel A). The relationship between this species and the FeS cluster was ruled out as we did not observe it with the FDP_E_Cter construct, either upon stepwise reduction with sodium dithionite or along the redox titration (green dotted spectrum in Figure 4 panel A). This led us to attribute this species to the flavin cofactor (see below).

Subsequently, we attempted to characterize the nature of the isolated flavin, which extraction from the FDP_E was only possible by thermal denaturation. The UV-Visible spectrum in the “as isolated” state flavin of FDP_E (Figure 4 panel C, yellow spectrum) exhibits the characteristic peaks of the UV-Visible spectra of an oxidized flavin, with maxima at 450 nm and 330-370 nm, arising from the n --> π* and π --> π* transitions [26,27], indicating an unaltered isoalloxazine ring. However, when the spectrum of the flavin isolated from FDP_E was compared with the commercial FMN and with the FMN extracted from *E. coli* FDP (pink spectrum) it shows a deviation mainly in the 330-370 nm band. [26,27] Upon reduction with excess of sodium dithionite, a spectrum of a neutral (blue) semiquinone was obtained (Figure 4 panel C) with maxima at 400 nm, a shoulder at ≈ 420nm and a very broad band with a maximum at 560 nm as described before [20]. As for the FDP_E, the complete reduction of this sample was not possible, even upon the addition of a large excess of sodium dithionite.

The fluorescence properties of this molecule were also studied. The emission spectra of FDP_E, partially reduced FDP_E (containing the “430 nm” species) and the flavin isolated from FDP_E (prepared as above) were almost identical with a single band with a maximum at 535 nm using both λ_Ex_= 455 nm (Figure S3 A) and λ_Ex_= 430 nm (FigureS3 B), whereas the excitation spectra (following the λ_Em_=520 nm, Figure S3 C) of these samples, presented broad bands at 460 nm and 360 nm, characteristic of oxidized FMN [26,27]. These similarity may indicate that the chemical nature of these species is identical among themselves and to other flavins.

Reverse phase HPLC revealed that this molecule has a higher hydrophobicity than the FMN, FAD or the riboflavin used as standards, and suggest heterogeneity of the extracted cofactor, as three elution peaks were observed, making impossible its direct identification by this technique (Figure S4).

### FeS cluster properties

The presence of an FeS cluster as part of a flavodiiron protein constitute a feature restricted to this class of FDPs and was characterized, by the first time, in the present work. As already mentioned in the amino acids sequence analysis’ section, the C-terminal FeS domain shares homology with a protein family “Fer4_19” that may harbour either a [3Fe-4S]^1+/0^ or a [4Fe-4S]^2+/1+^ cluster.

In Figure 5 panel A the EPR spectrum at 12 K of the as purified (oxidized) form of FDP_E (blue spectrum) showed a single species with a signal reminiscent of those of a [3Fe-4S]^1+^ cluster with a rhombic EPR signal with g-values very close to 2: 2.030, 2.019 and 2.003 (Table 1); the signal is observable up to 30 K, temperature at which it starts broadening, again a characteristic of oxidized [3Fe-4S] centres. Also, because of the presence of the flavin, we observed the contribution of a radical species with g ≍ 2, that must arise from its semiquinone form. The reduction of FDP_E with sodium dithionite in the presence of methyl viologen (Figure 5 panel A) completely bleached the [3Fe-4S]^1+^ EPR signal, being observed only a radical species signal with an isotropic signal at g ≍ 2, probably related with the presence of methyl viologen radical in the sample. The spectrum of FDP_E_Cter is identical to the one of the intact FDP_E acquired in the same conditions (Figure 5 panel A, pink spectra), which points to the presence of the same species in both proteins, the [3Fe-4S]^1+/0^ centre. The absence of an EPR signal in the reduced FDP_E and FDP_E_Cter samples indicated that there was no [4F-e4S]^2+/1+^ centre in the sample or that this centre would have a reduction potential lower than that of methyl viologen (-440 mV at pH 7).

**Table 1.** g-values for the EPR signals of the [3Fe-4S]^1+/0^ cluster in the oxidized samples of the different protein variants, obtained by theoretically simulating the spectra using SpinCount [23].

| Protein | $g_{\text{max}}$ | $g_{\text{med}}$ | $g_{\text{min}}$ |
| --- | --- | --- | --- |
| FDP_E | 2.003 | 2.019 | 2.030 |
| E_H441A | 2.003 | 2.017 | 2.030 |
| E_H441C | 2.001 | 2.016 | 2.031 |
| E_Cter | 1.999 | 2.015 | 2.028 |

**Figure 5.**
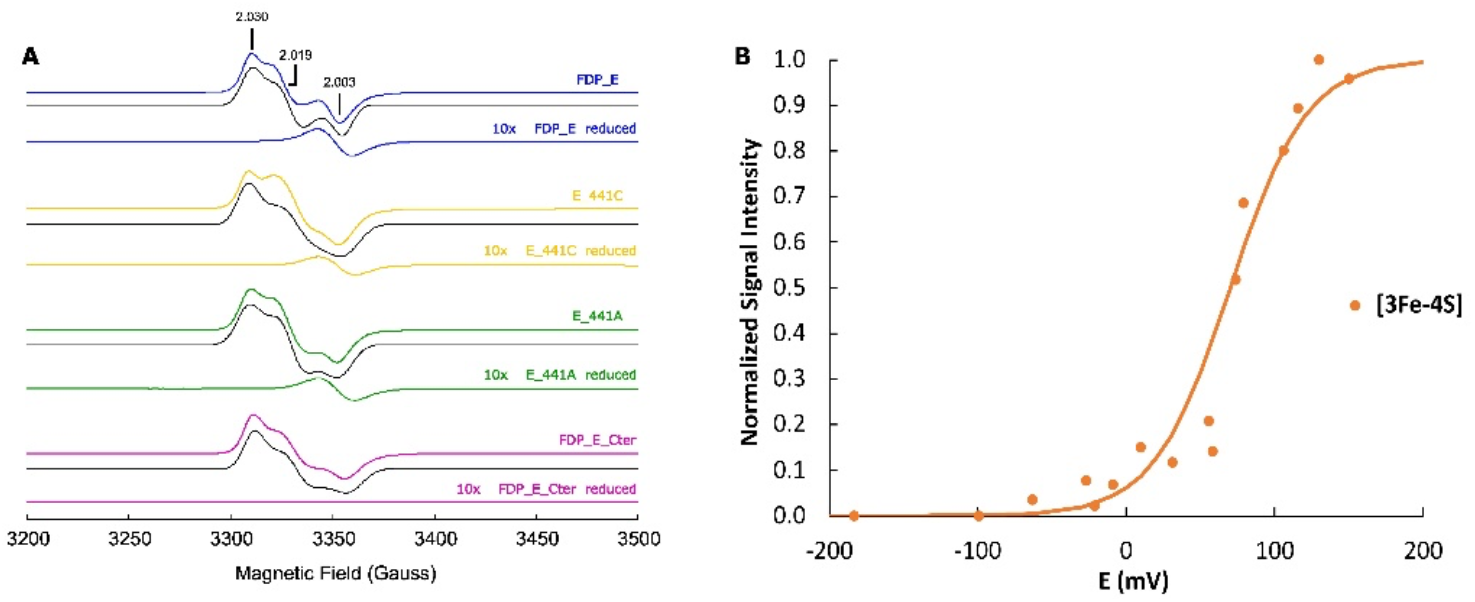
**(A) EPR spectra of 150 µM of FDP_E** (blue), **E_H441C** (yellow), **E_441A** (green) **and FDP_E_Cter** (pink) **in the oxidized and reduced** (spectra 10x multiplied) **forms**. The black line corresponds to the simulated spectra. **(B) Redox titration of the FeS cluster of FDP_E**. Temperature, 7 K; Microwave frequency, 9.39 GHz; Modulation amplitude, 1.0 mT; Microwave power, 2 mW. Spectral simulations and image preparation were performed using SpinCount [23].

To rule out the hypothesis that the FeS in FDP_E was originally a [4Fe4S]^2+/1+^ cluster, coordinated by the conserved cysteines and the histidine from the “Fer4_19” motif, that later undergoes degradation during the purification and manipulation procedures, giving rise to the 3Fe centre observed by EPR, we produced and purified two site-directed mutants of FDP_E. In these mutants the conserved histidine (residue 441) was replaced either by a cysteine (E_H441C) or by an alanine (E_H441A) to evaluate the possible role of the conserved histidine as a putative fourth ligand of the FeS cluster.

The EPR spectra of E_H441A and E_H441C variants in the “as purified” samples (Figure 5 green and yellow spectra respectively) revealed EPR signals very similar to the one of the wild type protein (Table 1). As for the wild type protein, the reduction with sodium dithionite and methyl viologen led to a bleaching of the EPR spectra, with no indication for the presence of a tetranuclear cluster, being only observed the radical species at g ≈ 2.

The presence of the [3Fe-4S]^1+/0^ cluster as a single species in FDP_E, E_H441A and E_H441C variants and FDP_E_Cter was verified also by Resonance Raman (RR) spectroscopy. The spectra of all samples exhibit a band centred at 345 cm^−1^ (Figure 6), indicative of FeS bridging vibrational mode of an oxidized 3Fe cluster. Note that no other bands were detected, apart from ice lattice modes [28], also present in the spectrum of the buffer (Figure S5). The spectra lack the characteristic pattern of [3F-e4S]^1+/0^ clusters, particularly the terminal FeS vibrational modes, derived from cysteinyl S ligands (2^nd^ most intense bands typically found at ca. 366 cm^−1^)[29–32]. Despite the incomplete assignment of characteristic RR features for a [3Fe-4S] ^1+/0^ cluster the detected signal is not attributable to binuclear or tetranuclear centres. The E_H441A and E_H441C variants have identical spectra to the wild type and C-terminal domain proteins indicating that the histidine residue cannot act as a non-cysteinyl ligand to the cluster and that its substitution by cysteine also does not lead to a [4Fe- 4S]^2+/1+^configuration. These results support that FDP_E houses a [3Fe-4S]^1+/0^ cluster, which agrees with the finding that Class E FDPs bear only three cysteines in the proposed cluster binding motif located in the C- terminal domain (CxHxxxCx_33_CP).

**Figure 6.**
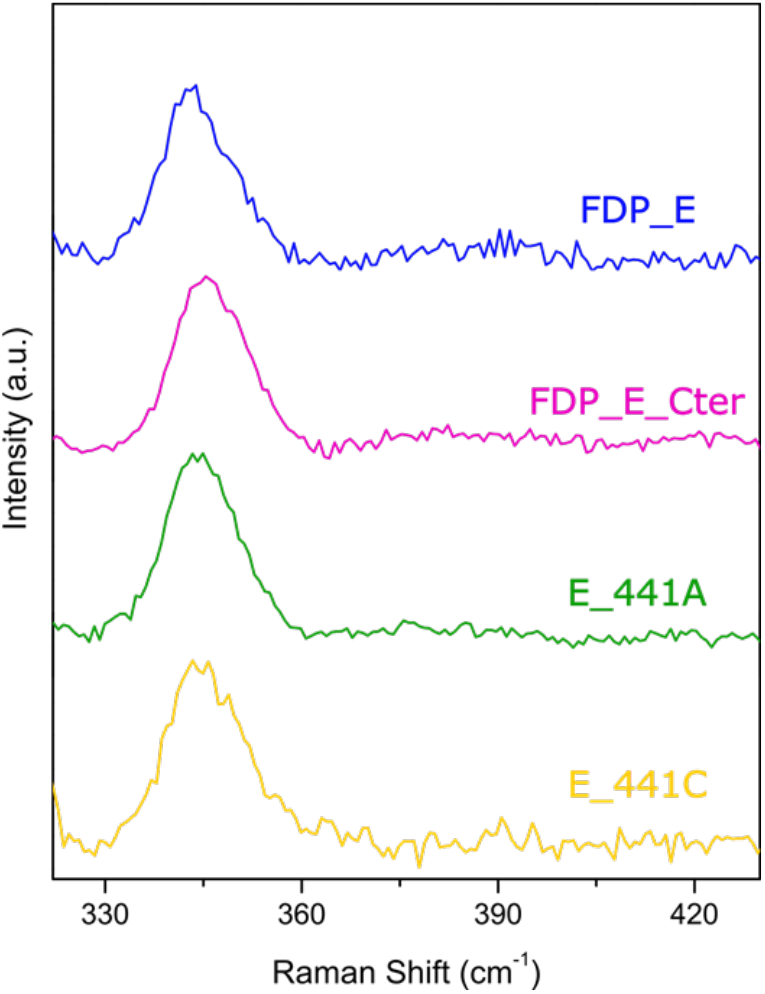
**Resonance Raman** spectra of FDP-E wild type (blue), **C-terminal domain** (pink) **and variants E_H441A** (green) **and E_H441C** (yellow) in 20 mM Tris-HCl pH 7.5, measured with 405 nm excitation.

The FDP_E was also anaerobically titrated and the EPR signal at g ≈2 was used to determine the cluster’s reduction potential. The experimental points were manually adjusted to aNernst equation for a monoelectronic transition, yielding a potential of 70 ± 10 mV (Figure 5 panel B). This potential is more positive when compared with the generality of the reduction potentials described for this type of centres, normally varying between –100 and –400 mV [5,33], but is similar, for example, to those reported for the [3Fe-4S]^1+/0^ centres of several succinate:quinone oxidoreductases (varying between +60 and +140 mV) [34,35].

### FDP_E electron transfer partner

The bioinformatic analysis of the *S. wolfei* genome did not reveal any canonical physiological electron donor for the FDPs (such as NADH:oxidoreductases or rubredoxins) [36]. Therefore, in an attempt to perform a kinetic characterization of FDP_E, we tested its reduction with the electron donors from other FDP’s classes: the *S. wolfei* FDP_H_Cter [9], the *C. difficile* FDP_F_Cter [24] and the *E. coli* flavorubredoxin reductase (NADH:flavorubredoxin oxidoreductase, NROR) [20] together with its rubredoxin domain, which has proved to be efficient for other FDPs whose electron donors are not known [37]. However, none of these systems was able to transfer electrons to FDP_E (data not shown). In addition, and given the structural homology of the C-terminal domain of FDP_E with ferredoxins (see “Amino acid sequence analysis and structural predictions” section) we also tested a commercially available Ferredoxin NADP^+^ Reductase, but also without success. Therefore, we could not determine whether this enzyme catalyses the reduction of any of the regular FDP substrates such as oxygen or nitric oxide.

## Conclusion

In summary, this work contributes to the characterization, by the first time, of one member of the class E FDPs. This protein is encoded in the genome of the anaerobic bacterium *Syntrophomonas wolfei subsp*. *wolfei* str. Goettingen G311 and by spectroscopic techniques was verified the presence of the three *in silico* predicted domains: the metallo-β-lactamase and the flavodoxin-like domains that constitute the typical FDP core and the C-terminal domain which containing a [3Fe- 4S]^1+/0^ centre.

Although the spectroscopic features of the metallo-β-lactamase like domain are in line with the features already described for the analogous domains in the other FDPs, the features of the flavin of the core domain are quite unusual.

The results obtained in the characterization of the flavodoxin domain points to the presence of a flavin moiety covalently bound to the polypeptide chain, as it is not extractable under mild conditions. Furthermore, a species never previously reported, to the best of our knowledge, has an UV-Visible spectrum with a pronounced peak at 430 nm. Moreover, the more unexpected feature of FDP_E flavin is the lower HPLC retention time in comparison with FMN, FAD and riboflavin molecules pointing either for the presence of a flavin molecule with some type of modification or to its degradation upon thermal denaturation of FDP_E, as suggested from the several peaks detected by reverse phase HPLC.

The characterization of the C-terminal domain of FDP_E revealed, for the first time in this family of proteins, the presence of a [3Fe-4S]^+1/0^ centre. This domain showed structural and spectroscopic similarities with [3F-e4S]^1+/0^ containing ferredoxins, suggesting the role of these centres as key players in the establishment of a new type of electron transfer centre from the C-terminal domain of FDP_E to the core domains.

## Materials and Methods

### Protein Expression and Anaerobic Purification

*Syntrophomonas wolfei subsp*. *wolfei* str. Goettingen G311 Swol_2363, encoding a class E FDP, was retrieved from NCBI and synthetized at GenScript Inc. USA. A truncated form comprising the C-terminal domain (residues 401 to 508) was also produced. The recombinant proteins were denominated as FDP_E and FDP_E_Cter, respectively. Additionally, two site-directed mutants were produced where the histidine (residue 441) localized in the conserved motif CXHX_3_C-X_32_CP presumably involved in the binding of a FeS cluster were substituted by an alanine (named as E_H441A) or a cysteine (named as E_H441C). In all syntheses were used codons optimized for expression in *E. coli* and the genes were cloned in a pET24a(+) vector, using NdeI and HindIII restriction sites. The plasmids were used to transform *E*. *coli* BL21(DE3) Gold competent cells (Nzytech). 300 mL of Luria Bertani medium containing 50 μg/mL kanamycin was inoculated with a single colony of *E. coli* BL21(DE3) Gold cells containing pET24a(+) FDP_E, FDP_E_Cter, E_H441A or E_H441C plasmids. Upon 16 h at 37 °C and 150 rpm, 60 mL of cell cultures of FDP_E, E_H441A and E_H441C was used to inoculate 2 L of M9 minimal medium, in 2 L Erlenmeyer flasks, containing 50 μg/mL kanamycin and supplemented with 0.1 mM of FeCl_2_. These cultures were grown at 37 °C and 160 rpm, until an optical density at 600 nm of approximately 0.4. At this point, 0.1 mM of FeCl_2_ was added and protein expression was induced with 100 μM of IPTG; cells were grown for additional 16 h at 25°C and 120 rpm. The expression of FDP_E_Cter was performed differently: 30 mL of pre inoculum culture was used to inoculate 1 L of LB medium, in 2 L Erlenmeyer flasks, containing 50 μg/mL kanamycin and 0.1 mM of FeCl_2._ This culture was grown at 37 °C and 150 rpm until an optical density at 600 nm of 1.2. At this point, the protein expression was induced with 500 μM of IPTG and a new addition of 0.1 mM FeCl_2_ was made; cells were grown for more 16 h at 25°C and 120 rpm. All the cells were harvested by centrifugation (7 000 g 10 min) resuspended in 20 mM Tris-HCl buffer pH 7.5 and disrupted by 3 cycles in a French-Press apparatus at 16000 psi (Thermo) in the presence of DNAse (Applichem). From this step forward all the purification procedures were realized under anaerobic conditions inside an anaerobic chamber (Coy Lab Products). We purified the holoenzyme (wild type and mutants) and the C-ter construct anaerobically, to avoid the eventual degradation of a [4Fe-4S]^2+/1+^ if it were present, as this type of clusters are frequently oxygen sensitive. We found no differences between the proteins purified aerobic- or anaerobically. The crude extracts were centrifuged at 25 000 g for 30 min and at 138 000 g for 2 h at 4 °C to remove cell debris and membrane fraction, respectively. The supernatants were maintained overnight at 4 °C anaerobically in an anaerobic encapsulated flask. The soluble extracts were loaded onto a Q-Sepharose Fast-Flow column (65 mL, GE Healthcare) previously equilibrated with 20 mM Tris-HCl buffer pH 7.5 containing 18 % glycerol (buffer A). The proteins were eluted with a linear gradient from buffer A to 20 mM Tris-HCl pH 7.5 containing 18 % glycerol and 500 mM NaCl. The eluted fractions were analyzed by 15 % SDS-PAGE and UV-Visible spectroscopy. Fractions with the desired protein were pooled and concentrated. The selected fractions were loaded into a Fractogel column (20 mL, Merck Millipore) previously equilibrated with buffer A and eluted with a linear gradient from A to 20 mM Tris-HCl pH 7.5 containing 18 % glycerol and 400 mM NaCl. The FDP_E_Cter was subjected to a third chromatographic step on a size exclusion S200 column (120 mL, GE Healthcare) equilibrated with 20 mM Tris–HCl, pH 7.5 containing 18 % glycerol and 150 mM NaCl. Fractions containing pure proteins were pooled and concentrated.

### Protein, metal and flavin quantification

All the proteins (FDP_E, FDP_E_Cter, E_H441A and E_H441C) were quantified using the Bicinchoninic Acid Kit (Thermo) and bovine serum albumin as standard. Iron content was determined by inductively coupled plasma-atomic emission spectroscopy (ICP-AES) at Laboratório de Análises, Chemistry Department, CQFB/REQUIMTE, FCT-UNL). For flavin quantification, an FDP_E was incubated for 45 min at 95 °C, centrifuged at 16 000 g for 10 min and the supernatants were used to measure the absorbance spectrum; the flavin content was quantified using ɛ_446nm_ = 12.5 mM^−1^.cm^−1^ [27]. The flavin type was analyzed by reverse phase HPLC at the ITQB Small Molecule Analysis facility, as previously described [24].

### Protein quaternary structure determination

The quaternary structures were determined by size exclusion chromatography using a 25 mL Superdex 200 10/300 GL Increase column (GE Healthcare) for FDP_E, as previously described in [9,24], and a 25 mL Superdex S75 10/300 GL column in the case of FDP_E_Cter. A mixture containing conalbumin (75 kDa), ovalbumin (44 kDa), carbonic anydrase (29 kDa), ribonuclease a (13.7 kDa), aprotinin (6.5 kDa), and dextran blue (2000 kDa) as a void volume marker was used as the standard in the Superdex S-75 10/300 GL column.

### UV-Visible spectroscopy

FDP_E and FDP_E_Cter samples were prepared inside an anaerobic chamber (Coy Lab Products) to a final concentration of 30 μM in 50 mM Tris-HCl pH 7.5 containing 18% and in the presence of an O_2_ scavening system composed by: 10 mM glucose, 375 nM glucose oxidase and 750 nM catalase. Once the UV-Visible initial spectra (corresponding to the oxidized state) were obtained the samples were reduced by sequential additions of a buffered sodium dithionite solution (50 mM Tris-HCl pH 9).

The reduction potential of FDP_E flavin was determined by UV-Visible spectroscopy, using a Shimadzu UV-1603 spectrophotometer. 30 μM of FDP_E in 50 mM Tris–HCl, pH 7.5 containing 18% glycerol and in the presence of the O_2_ scavenging system (above mentioned) was anaerobically titrated by consecutive additions of a buffered solution of a sodium dithionite solution (50 mM Tris-HCl pH 9) in the presence of a mixture of redox mediators (1 μM of each, assay final concentration): potassium ferricyanide (E’_0_= +430mV), N,N dimethyl-p-fenilenodiamine (E’_0_= +340mV), 1,2- naphtoquinone-4-sulphonic acid (E’_0_= +215 mV), 1,2 naphtoquinone (E’_0_= +180 mV), trimethylhydroquinone (E’_0_= +115 mV), 1,4 naphtoquinone (E’_0_= +60 mV), 5-hydroxy-1,4-naphthoquinone (E’_0_ = +30 mV), duroquinone (E’_0_= +5 mV), menadione (E’_0_= 0 mV), plumbagin (E’_0_= -40 mV), resorufin (E’_0_= -51 mV), indigo trisulphonate (E’_0_= -70 mV), phenazine (E’_0_= -125 mV), 2-hydroxy-1,4-naphthoquinone (E’_0_= - 152 mV), anthraquinone-2-sulphonate (E’_0_= -225 mV), phenosafranine (E’_0_= -255 mV), safranine (E’_0_= -280 mV) and neutral red (E’_0_= -325 mV).

A combined Pt electrode (Ag/AgCl in 3.5 M KCl, as reference) was used and calibrated at 20 °C against a saturated quinhydrone solution (pH 7). The reduction potentials were calculated in relation to the standard hydrogen electrode. The experimental points represented are the result of three independent redox titrations. The experimental data was manually adjusted with a Nernst equation.

### Electron paramagnetic resonance (EPR) spectroscopy

Electron paramagnetic resonance (EPR) spectroscopy was performed using a Bruker EMX spectrometer equipped with an Oxford Instruments ESR-900 continuous-flow helium cryostat and a perpendicular mode rectangular cavity.

The as isolated protein samples were prepared anerobically to final concentrations of 150 µM (for FDP_E, E_H441A and E_H441C) or 100 µM (in the case of FDP_E_Cter). A reduced sample for all the proteins was prepared anaerobically by incubation with sodium dithionite (10x excess) and methyl viologen (5x excess). EPR spectra were obtained at 12 K and were simulated using the program SpinCount [23]

The reduction potentials of the diiron and the FeS centres were determined by EPR spectroscopy. 100 µM of FDP_E samples were prepared under anaerobic conditions in 50 mM Tris–HCl, pH 7.5 containing 18% glycerol and in the presence of the O_2_ scavening system. These samples were titrated by consecutive additions of a buffered sodium dithionite solution (50 mM Tris-HCl pH 9) in the presence of a mixture of redox mediators (30 μM of each, assay final concentration) similar to the one mentioned for UV-Visible spectroscopy with the addition of two mediators: phenazine methosulfate (E’_0_= +80mV) and methylene blue (E’_0_= +11mV). The electrode and the respective calibration are the same as mentioned in the above section. The experimental points represented correspond to the normalized intensity of the signal at g =1.85 for the diiron and g ≈ 2 for the FeS centres and result from two independent redox titrations (independent normalization of the two datasets). The experimental data was manually adjusted to the appropriate Nernst equations: monoelectronic Nernst equation for the FeS centre, and a Nernst equation for two consecutive monoelectronic transitions for the diiron centre.

### Ressonance Raman spectroscopy

Resonance Raman (RR) spectra were acquired with a Raman spectrometer (Horiba LabRam HR- 800) equipped with a 1200 lines/mm grating and a liquid-nitrogen-cooled CCD detector. An Olympus 20x objective was used for laser focusing onto the sample and light collection in the backscattering geometry. Spectra of FDP_E, FDP_E_Cter, E_H441A and E_H441C variants were measured using a 405 nm diode laser (Toptica Photonics AG) at −190 °C, from 3 μL aliquots of the samples (0.2–0.5 mM) placed in a microscope stage (Linkham THMS 600). Experiments were performed with a laser power of 9 mW and 40- 60s accumulation times. Up to 16 spectra were co-added in each measurement to improve the signal-to-noise ratio (S/N).

### Fluorescence spectroscopy

The flavin molecule isolated from FDP_E was analysed by steady-state fluorescence in a Varian Cary Spectrometer. Sample concentration was 10 μM to avoid saturation of the emission spectra. Excitation spectrum was recorded between 250 and 500 nm at λ_Em_= 520 nm, whereas the emission spectrum was recorded between 500 and 800 nm with excitation at λ_Exc_= 455 nm and λ_Exc_= 430 nm, with a slit width of 10 nm in both cases, in order to excite the flavin molecule.

## Author Contributions

The project was coordinated and designed by MT and FF. FF and MCM performed the experiments and analysed the data. The Raman Spectroscopic studies were realized and analysed by CS. All authors contributed to the manuscript writing and approved the final version of the article.

## References

1. Martins MC, Romão C V, Folgosa F, Borges PT, Frazão C & Teixeira M (2019) How superoxide reductases and flavodiiron proteins combat oxidative stress in anaerobes. Free Radic Biol Med 140, 36–60.

2. Wasserfallen A, Ragettli S, Jouanneau Y & Leisinger T (1998) A family of flavoproteins in the domains archaea and bacteria. Eur J Biochem 254, 325–332.

3. Folgosa F, Martins MC & Teixeira M (2018) Diversity and complexity of flavodiiron NO/O2 reductases. FEMS Microbiol Lett 365.

4. Py B & Barras F (2010) Building Feg-S proteins: Bacterial strategies. Nat Rev Microbiol 8, 436–446.

5. Liu J, Chakraborty S, Hosseinzadeh P, Yu Y, Tian S, Petrik I, Bhagi A & Lu Y (2014) Metalloproteins containing cytochrome, iron-sulfur, or copper redox centers. Chem Rev 114, 4366–4369.

6. Crack JC, Green J, Thomson AJ & Brun NEL (2014) Iron-sulfur clusters as biological sensors: The chemistry of reactions with molecular oxygen and nitric oxide. Acc Chem Res 47, 3196–3205.

7. Lukianova OA & David SS (2005) A role for iron-sulfur clusters in DNA repair. Curr Opin Chem Biol 9, 145–151.

8. Meyer J (2008) Iron-sulfur protein folds, iron-sulfur chemistry, and evolution. Journal of Biological Inorganic Chemistry 13, 157–170.

9. Martins MC, Alves CM, Teixeira M & Folgosa F (2023) The flavodiiron protein from *Syntrophomonas wolfei* has five domains and acts both as an NADH:O2 or an NADH:H2O2 oxidoreductase. FEBS Journal, 291, 1275–1294.

10. Dementin S, Burlat B, Fourmond V, Leroux F, Liebgott PP, Abou Hamdan A, Leéger C, Rousset M, Guigliarelli B & Bertrand P (2011) Rates of intra- and intermolecular electron transfers in hydrogenase deduced from steady-state activity measurements. J Am Chem Soc 133, 10211–10221.

11. Jumper J, Evans R, Pritzel A, Green T, Figurnov M, Ronneberger O, Tunyasuvunakool K, Bates R, Žídek A, Potapenko A, Bridgland A, Meyer C, Kohl SAA, Ballard AJ, Cowie A, Romera-Paredes B, Nikolov S, Jain R, Adler J, Back T, Petersen S, Reiman D, Clancy E, Zielinski M, Steinegger M, Pacholska M, Berghammer T, Bodenstein S, Silver D, Vinyals O, Senior AW, Kavukcuoglu K, Kohli P & Hassabis D (2021) Highly accurate protein structure prediction with AlphaFold. Nature 596, 583– 589.

12. Varadi M, Anyango S, Deshpande M, Nair S, Natassia C, Yordanova G, Yuan D, Stroe O, Wood G, Laydon A, Zídek A, Green T, Tunyasuvunakool K, Petersen S, Jumper J, Clancy E, Green R, Vora A, Lutfi M, Figurnov M, Cowie A, Hobbs N, Kohli P, Kleywegt G, Birney E, Hassabis D & Velankar S (2022) AlphaFold Protein Structure Database: Massively expanding the structural coverage of protein-sequence space with high-accuracy models. Nucleic Acids Res 50, D439–D444.

13. Romão C V., Vicente JB, Borges PT, Victor BL, Lamosa P, Silva E, Pereira L, Bandeiras TM, Soares CM, Carrondo MA, Turner D, Teixeira M & Frazão C (2016) Structure of *Escherichia coli* Flavodiiron Nitric Oxide Reductase. J Mol Biol 428, 4686–4707.

14. Abdulaziz EN, Bell TA, Rashid B, Heacock ML, Begic T, Skinner OS, Yaseen MA, Chao LH, Mootha VK, Pierik AJ & Cracan V (2022) A natural fusion of flavodiiron, rubredoxin, and rubredoxin oxidoreductase domains is a self-sufficient water-forming oxidase of *Trichomonas vaginalis*. Journal of Biological Chemistry 298(8):102210.

15. Kelley LA, Mezulis S, Yates CM, Wass MN & Sternberg MJE (2015) The Phyre2 web portal for protein modeling, prediction and analysis. Nat Protoc 10, 845–858.

16. Bruschi M & Guerlesquin F (1988) Structure, function and evolution of bacterial ferredoxins. FEMS Microbiol Lett 54, 155–175.

17. Gilep A, Varaksa T, Bukhdruker S, Kavaleuski A, Ryzhykau Y, Smolskaya S, Sushko T, Tsumoto K, Grabovec I, Kapranov I, Okhrimenko I, Marin E, Shevtsov M, Mishin A, Kovalev K, Kuklin A, Gordeliy V, Kaluzhskiy L, Gnedenko O, Yablokov E, Ivanov A, Borshchevskiy V & Strushkevich N (2023) Structural insights into 3Fe–4S ferredoxins diversity in *M. tuberculosis* highlighted by a first redox complex with P450. Front Mol Biosci 9.

18. Bak DW (2006) A study of the CDGSH protein family: biophysical and bioinformatic analysis of the [2fe-2s] cluster protein mitoneet. PhD Thesis, Boston University.

19. Frazão C, Silva G, Gomes CM, Matias P, Coelho R, Sieker L, Macedo S, Liu MY, Oliveira S, Teixeira M, Xavier A V, Rodrigues-Pousada C, Carrondo MA & Le Gall J (2000) Structure of a dioxygen reduction enzyme from *Desulfovibrio gigas*. Nat Struct Biol. 7(11):1041–1045.

20. Vicente JB & Teixeira M (2005) Redox and spectroscopic properties of the *Escherichia coli* nitric oxide-detoxifying system involving flavorubredoxin and its NADH-oxidizing redox partner. Journal of Biological Chemistry 280, 34599–34608.

21. Vicente JB, Tran V, Pinto L, Teixeira M & Singha U (2012) A detoxifying oxygen reductase in the anaerobic protozoan *Entamoeba histolytica*. Eukaryot Cell 11, 1112– 1118.

22. Vicente JB, Testa F, Mastronicola D, Forte E, Sarti P, Teixeira M & Giuffrè A (2009) Redox properties of the oxygen-detoxifying flavodiiron protein from the human parasite *Giardia intestinalis*. Arch Biochem Biophys 488, 9–13.

23. 23. Petasis DT & Hendrich MP (2015) Quantitative interpretation of multifrequency multimode EPR spectra of metal containing proteins, enzymes, and biomimetic complexes. Methods in Enzymology- Academic Press Inc 563, 171–208.

24. Folgosa F, Martins MC & Teixeira M (2018) The multidomain flavodiiron protein from *Clostridium difficile* 630 is an NADH:oxygen oxidoreductase. Sci Rep 8, ,10164.

25. Vicente JB, Carrondo MA, Teixeira M & Frazão C (2008) Structural Studies on Flavodiiron Proteins. Methods in Enzymology 437, 3–19.

26. Weber S & Schleicher E (2014) Flavins and Flavoproteins - Methods and Protocols. Methods in Molecular Biology Human Press Inc 131, 191–228.

27. Chapman SK & Reid GA (1999) Flavoprotein Protocols. Methods in Molecular Biology Human Press Inc 131, 1–48.

28. Fu W, O’Handley S, Cunningham RP & Johnson MK (1992) The role of the iron-sulfur cluster in *Escherichia coli* endonuclease III: A resonance Raman study. Journal of Biological Chemistry 267, 16135–16137.

29. Todorovic S & Teixeira M (2018) Resonance Raman spectroscopy of Fe–S proteins and their redox properties. Journal of Biological Inorganic Chemistry 23, 647–661.

30. Todorovic S, Leal SS, Salgueiro CA, Zebger I, Hildebrandt P, Murgida DH & Gomes CM (2007) A spectroscopic study of the temperature induced modifications on ferredoxin folding and iron-sulfur moieties. Biochemistry 46, 10733–10738.

31. Caserta G, Zuccarello L, Barbosa C, Silveira CM, Moe E, Katz S, Hildebrandt P, Zebger I & Todorovic S (2022) Unusual structures and unknown roles of FeS clusters in metalloenzymes seen from a resonance Raman spectroscopic perspective. Coord Chem Rev 452.

32. Spiro TG & ROMAN S. Czernoszewlcz (1993) Resonance Raman Spectroscopy of Metalloproteins. Methods in Enzymology 246.

33. Moura JJ, Macedo AL, Goodfellow BJ & Moura I (2004) Ferredoxins Containing One [3Fe–4S] Cluster. *Desulfovibrio Gigas* Ferredoxin II–Solution Structure. *Handbook of Metalloproteins*, Wiley.

34. Hägerhäll C, Aasa R, Von Wachenfeldt C & Hederstedt L (1992) Two Hemes in *Bacillus subtilis* succinate:menaquinone oxidoreductase (complex II). Biochemistry 31, 7411–7421.

35. Fernandes AS, Pereira MM & Teixeira M (2001) The Succinate Dehydrogenase from the thermohalophilic bacterium *Rhodothermus marinus*: Redox-Bohr Effect on Heme bL. J Bioenerg Biomembr 33, 343–352.

36. Sieber JR, Sims DR, Han C, Kim E, Lykidis A, Lapidus AL, McDonnald E, Rohlin L, Culley DE, Gunsalus R & McInerney MJ (2010) The genome of *Syntrophomonas wolfei:* New insights into syntrophic metabolism and biohydrogen production. Environ Microbiol 12, 2289–2301.

37. Kint N, Feliciano CA, Martins MC, Morvan C, Fernandes SF, Folgosa F, Dupuy B, Texeira M & Martin-Verstraete I (2020) How the Anaerobic Enteropathogen *Clostridioides difficile* Tolerates Low O2 Tensions. mBio. 11(5),e01559–20.

