## Supplemental Material for "An iron-sulfur cluster as a new metal centre in a flavodiiron protein"

### Supporting Information

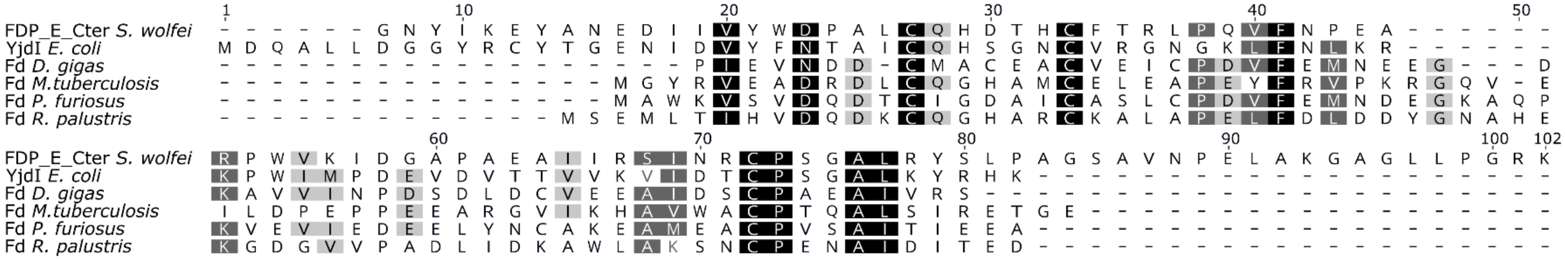


**Figure S1. Amino acid sequences alignment of FDP_E_Cter from *S. wolfei* with YjdI from *E. coli* and [3Fe-4S]$^{1+/0}$ ferredoxins from *D. gigas*, *M. tuberculosis*, *P. furiosus* and *R. palustris*.**

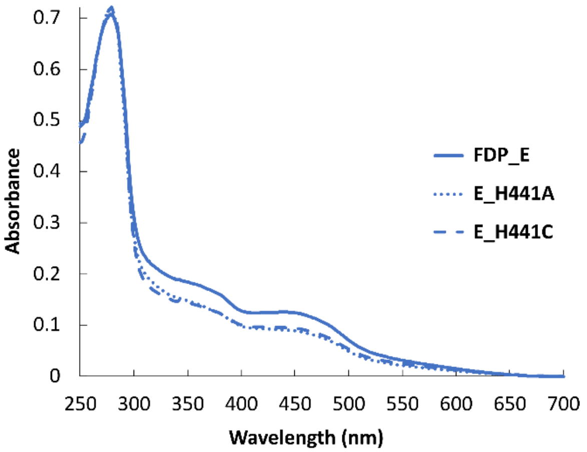


**Figure S2. UV-Visible spectra of 18 µM of FDP_E, E_H441A and E_H441C** in 50 mM Tris-HCl pH 7.5 containing 18 % glycerol.

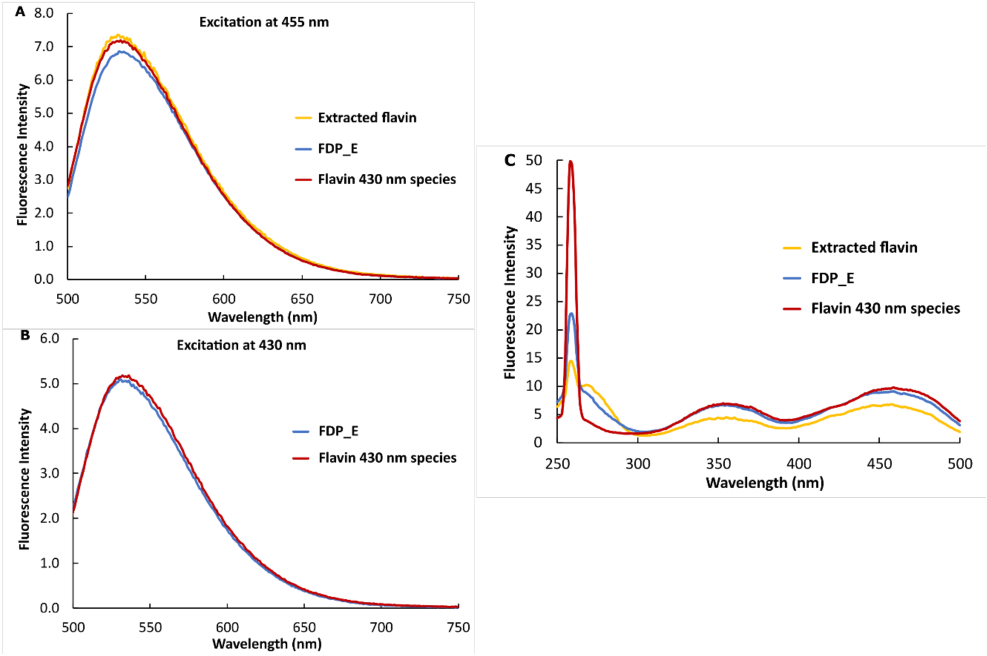


**Figure S3. Fluorescence spectra of FDP_E, the respective flavin obtained after denaturation and the partially reduced FDP_E containing the "430 nm" species. (A)** Emission spectra with $\lambda_{Ex}$= 455 nm. **(B)** Emission spectra with $\lambda_{Ex}$= 430 nm. **(C)** Excitation spectra monitoring the $\lambda_{Em}$= 520 nm.

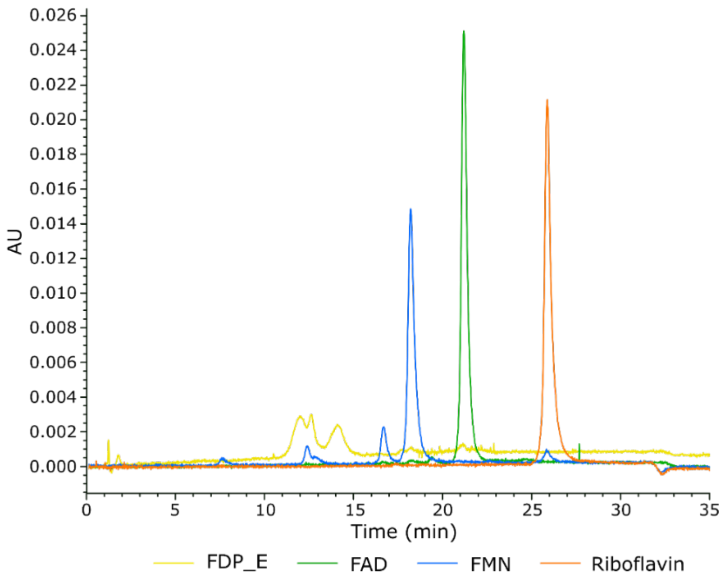


**Figure S4. Chromatogram of reverse phase HPLC of the extracted flavine obtained after the thermal denaturation of FDP_E and commercial FMN, FAD and Riboflavin.**

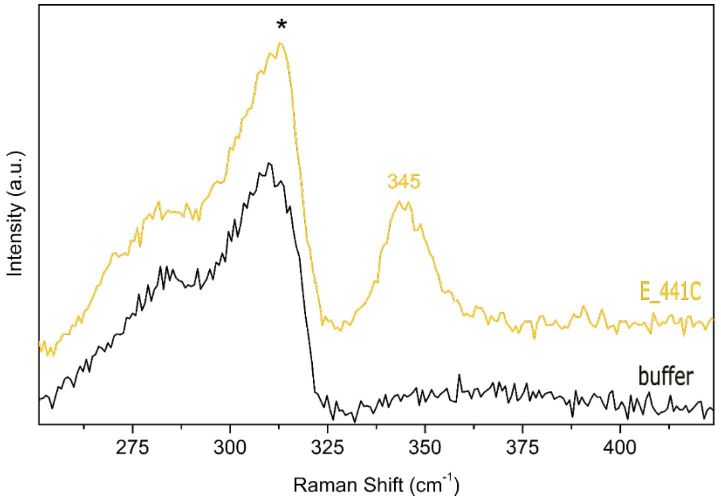


**Figure S5. Resonance Raman spectra of FDP variant E_H441C** (yellow) **and the 20 mM Tris-HCl pH 7.5 buffer** (black). Spectra were obtained with 405 nm excitation. The asterisk indicates the lattice modes of ice.
